# Upregulation of a nonsense mediated decay (NMD) insensitive CFTR mRNA isoform has therapeutic potential for the treatment of 3’ CFTR PTC variants

**DOI:** 10.1101/2024.07.01.601512

**Authors:** Normand E. Allaire, Mathew Armstrong, Jae Seok Yoon, Priyanka Bhatt, Jan Harrington, Yi Cheng, Hillary Valley, Caitlin Macadino, Hermann Bihler, Andrey Sivachenko, Martin Mense, Calvin Cotton

**Affiliations:** CFFT Lab, Cystic Fibrosis Foundation, Lexington, MA 02421, USA

**Keywords:** Cystic Fibrosis, CFTR, W1282X, mRNA isoform, mRNA processing, alternative polyadenylation, antisense oligonucleotide

## Abstract

**Background:** Nonsense or Premature Termination Codon (PTC) mutations of the *CFTR* gene are pathogenic and found in ∼10% of North American people with cystic fibrosis. PTC mutations induce Nonsense-Mediated mRNA Decay (NMD), leading to a substantial (∼80-90%) reduction in full-length mRNA. This reduction is a key contributor to PTC mutation-related pathology. Various approaches to evade NMD and preserve the impacted mRNA transcript have been explored but have not progressed to clinical development, leaving NMD a significant hurdle for PTC readthrough therapy.

**Methods:** Antisense oligonucleotides (ASOs) were tiled across intron 22 splice donor (SD) and acceptor (SA) sites of the *CFTR* gene. Immortalized airway cells were treated with SD and SA ASOs, and those yielding the greatest increase in a *CFTR* NMD-insensitive mRNA isoform (e22 trunc mRNA) were tested in combinations. Top SD/SA ASO pairs were assessed for their impact on e22 trunc mRNA via ddPCR, e22 trunc protein via western blot, and CFTR-mediated chloride (Cl^-^) transport via transepithelial electrophysiological measurements in immortalized and primary human bronchial epithelial (hBE) cell cultures.

**Results:** We demonstrate that e22 trunc mRNA generates a truncated CFTR protein whose Cl^-^ transport function can be enhanced with elexacaftor/tezacaftor/ivacaftor (ETI) treatment. ASO and ETI treatment in combination restore ∼20% and 25% of wild-type CFTR Cl^-^ transport function in immortalized epithelial and primary hBE cells homozygous for *CFTR* W1282X, respectively.

**Conclusions:** This study lays the groundwork for advancing ASO-mediated upregulation of e22 trunc mRNA and protein as a therapeutic approach for cystic fibrosis caused by 3’-terminal *CFTR* PTC mutations.

## Background

Cystic fibrosis (CF) is a severe autosomal recessive disease caused by mutations in the gene encoding the cystic fibrosis transmembrane regulator (CFTR) protein. The most common pathological features are pancreatic insufficiency and bronchiectasis with chronic airway infections resulting in respiratory failure and premature death (1). Previously, only symptomatic treatments were available to people with CF (pwCF) such as pancreatic enzymes to aid digestion, airway clearance techniques to reduce mucus accumulation, and antibiotics for pulmonary infections (2). More recently, a new class of drugs, known as CFTR modulators, that target the basic molecular defect(s) to improve CFTR trafficking to the plasma membrane and/or channel gating have been employed therapeutically. As of April 2023, 804 variants have been reported on by the **C**linical and **F**unctional **Tr**anslation of CFTR (CFTR2) project (https://cftr2.org). 719 of these variants are classified as CF-causing. Based on genotype, ∼93-94% of pwCF could benefit from the therapies directed at the underlying cause of their disease (3,4); however, the remaining ∼6-7% have *CFTR* variants including nonsense mutations that are not approved for current modulator treatments.

Here we show the existence of a rare naturally occurring, alternatively polyadenylated (ApA) CFTR mRNA isoform that terminates in intron 22, which we term e22 trunc mRNA. Unlike full-length W1282X (FL-W1282X) mRNA, e22 trunc mRNA derived from the W1282X allele is not subject to NMD. Furthermore, ASO-mediated blockage of exon 22/23 splicing enhances intron 22 ApA usage and increases expression of this NMD insensitive mRNA in PTC variants downstream of exon 22. The resultant truncated protein product is trafficked to the plasma membrane and exhibits anion channel activity that can be ∼10-fold enhanced by treatment with a CFTR potentiator. This approach may provide clinical benefit for pwCF that carry certain 3’ PTC variants such as W1282X, Q1313X, E1371X, etc.

## Methods

### Cell culture

16HBE14o-parental cells, and CFF-16HBEge CFTR W1282X (referred to as 16HBEge W1282X), and *CFF-16HBEge CFTR delta exon 23-3’UTR* clonal lines (*16HBEge-ΔE23-3’UTR*) were grown as described previously (5). Fischer Rat Thyroid (FRT) cells (6) and primary hBE cells (7) were cultured as previously described. Intestinal organoids (IOs) were cultured as previously described (https://cdn.stemcell.com/media/files/pis/10000003510-PIS_08.pdf).

### Generation of gene edited cell lines

16HBE14o-parental cells were edited with CRISPR/cas9 and the resulting 16HBEge W1282X and *16HBEge-ΔE23-3’UTR* cells were cloned and cultured as previously described (5). The following targeting component of the crRNA guide RNAs were used to generate the exon 23 to 3’ UTR deletion clones: 5’-TGCTCAGTTATAGTATATAA-3’ and 5’-TTAGTTATCTGTTTAAACTA-3’.

### Free uptake ASO treatments of 16HBEge W1282X & 16HBE14o-

Steric blocking ASOs were designed and purchased from IDT (Coralville, Iowa) with Phosphorothioate (PS) backbone and 2-O-Methoxyethyl (2’-MOE) modifications at each nucleotide (**Supplementary Table S1**). Lyophilized steric blocking ASOs were resuspended in PBS +Mg^2+^+Ca^2+^ to 1 mM stock concentration and stored in frozen aliquots. 16HBEge W1282X and 16HBE14o-lines were seeded at 150,000 cells/well in 24 well plates and grown as previously described. After 24 h, media was refreshed with addition of ASO. RNA was harvested after 48 h of treatment.

### Free uptake ASO treatments in primary hBE cells

2’-MOE ASOs were resuspended as above and diluted into the cell culture media. Undifferentiated basal cells were treated with ASO-supplemented media for 48 h and resupplied with every media change throughout differentiation for 21 days.

### ddPCR Isoform quantification

RNA was isolated with the Aurum Total RNA Mini Kit (Bio-Rad). cDNA synthesis was performed with the iScript Advanced cDNA Synthesis Kit for RT-qPCR (Bio-Rad). Droplets were generated and quantitated on the QX200 Droplet Digital PCR System (Bio-Rad). Data was analyzed with Quantasoft software version 1.7.4. ddPCR probes designed to the junction of exon 25/26 were used as a proxy of full-length WT CFTR (FL-WT) or FL-W1282X mRNA and probes designed to exon 22/intron 22 junction including the first 140 bp of intron 22 were used as a proxy of exon 22 truncated CFTR mRNA **(Supplementary Table S2)**.

### Long read 3’ RACE analysis

Reverse transcription was completed using Oligo d(T)23 VN_T3 = 5’-GCAATTAACCCTCACTAAAGGTTTTTTTTTTTTTTTTTTTTTTTVN-3’ and ProtoScript® II First

Strand cDNA Synthesis Kit (New England Biolabs) according to manufacturers recommendations. PCR amplification with was Q5^®^ High-Fidelity DNA Polymerase (New England Biolabs) with CFTR Exon 8 forward primer (5’-TTTCTGTTGGTGCTGATATTGCCTCAGGGTTCTTTGTGGTGT-3’) and T3 reverse primer (5’-ACTTGCCTGTCGCTCTATCTTCGCAATTAACCCTCACTAAAGG-3’) according to manufacturer’s recommendations. Amplicons were barcoded using SQK-LSK109 (Nanopore), purified with AMPure (Beckman Coulter), and sequenced on Minion (Nanopore).

### Western blot analyses

Western blot analyses were performed as described previously (6), except the loading control, ACTB was detected with beta actin antibody C4 (sc-47778) from Santa Cruz Biotechnology. Blots were imaged using the ChemiDoc MP Imaging System (BioRad) or ChemiDoc XRS+ Gel Documentation System (BioRad) and analyzed with the accompanying image analysis software.

### Electrophysiology

Conductance (FRT cells) and equivalent current (I_eq_) studies (CF hBE at air liquid interface (ALI) and 16HBEge W1282X cells) with the TECC-24 system were performed as previously described (5,6,7).

### Statistics

Statistical analysis of ddPCR data was performed using Microsoft Excel (Microsoft Corporation). All data was presented as mean ± standard deviation (STD). Two-tailed unpaired *t*-test was used to determine statistical significance of data. ANOVA with Tukey post-hoc test was performed on TECC-24 l_eq_ and Conductance data. P-values were calculated using Origin version 10.0.0.154 (OriginLab Corporation).

## Results

### Intron 22 ApA usage results in a naturally occurring, lowly expressed exon 22 truncated CFTR mRNA isoform that is insensitive to NMD and encodes a truncated protein

A partial CFTR exon 22 truncated mRNA isoform was first identified by Burch (8) and subsequently other exon 22 truncated isoforms have been reported in the Ensembl database (ENST00000648260.1 & ENST00000649406.1). We also detected an exon 22 truncated mRNA isoform in 16HBE14o-cells using 3’ RACE with a 5’ primer targeting exon 8 followed by long read sequencing. As with the transcripts reported by Burch and in the Ensembl database, exon 22 truncated CFTR transcripts extend ∼140 bp into intron 22 which encodes for alternative 9 additional amino acid residues and 3’ UTR. The exon 22 truncated transcripts terminate with 10-30 post-transcriptionally added adenosine residues that align with putative consensus hexanucleotide ApA and dinucleotide cleavage sites indicating usage of an intron 22 ApA site (**Supplementary Figure S1**). These transcripts are referred to as “e22 trunc”. *CFTR* cDNA long read isoform sequencing with 5’ UTR and 3’ intron 22 targeting primers was used to confirm the presence of “complete” (containing exons 1-22) e22 trunc mRNA in 16HBE14o-cells (**Figure 1**).

**Figure 1.**
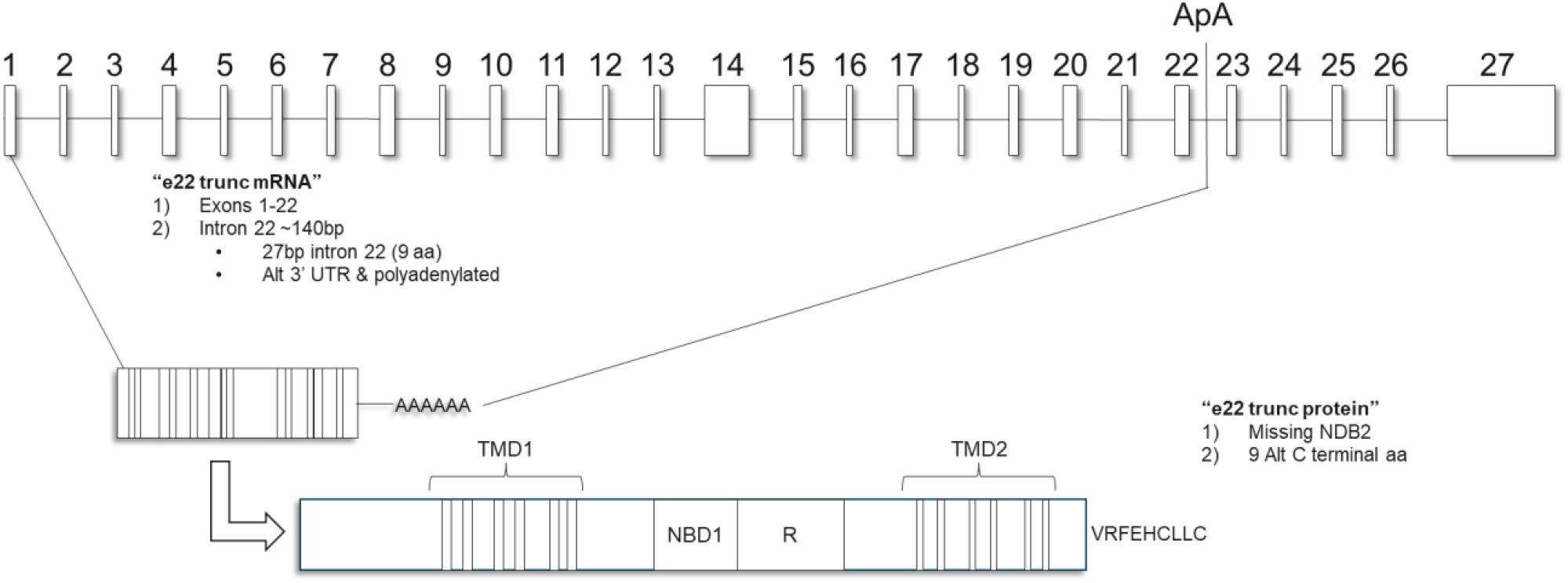
CFTR genomic locus, e22 trunc mRNA and protein. *CFTR* genomic locus with intron 22 ApA sites. Introns not shown to scale. e22 trunc mRNA representation showing alternative 3’UTR and poly A tail. e22 trunc protein representation showing additional 9 alternative amino acids and missing NBD2.

To our knowledge, this is the first report of exon 22 truncated CFTR mRNA containing all 22 upstream exons. Translation of e22 trunc mRNA yields a truncated CFTR protein (1239+9 aa) that retains transmembrane domains 1 and 2 (TMD1 & TMD2), nucleotide binding domain 1 (NBD1), R domain, and partial NBD2 followed by 9 intron 22-encoded alternative amino acid residues (VRFEHCLLC).

e22 trunc mRNA transcript derived from a *CFTR* W1282X allele should be insensitive to NMD. To evaluate this, FL-WT, FL-W1282X, and e22 trunc mRNA expression levels were quantified in hBEs, IOs, and immortalized airway cells with wild type (WT) or W1282X^+/+^ *CFTR* variant (**Supplementary Table S3**). As expected, FL-W1282X mRNA was reduced by ∼80-90% relative to FL-WT levels; whereas e22 trunc mRNA expression had similar low expression levels (8.4% ± 2.5) in both WT and *W1282X* cells suggesting that unlike FL-W1282X, e22 trunc mRNA is not subject to NMD.

### Therapeutic rationale for upregulation of a NMD insensitive CFTR mRNA isoform for the treatment of 3’ CFTR PTC variants

Since e22 trunc mRNA from W1282X allele should be insensitive to NMD, we sought to quantify the half-life (t_1/2_) of FL-WT, FL-W1282X, and e22 trunc mRNAs using actinomycin D time courses. As illustrated in **Figure 2A**, t_1/2_ of e22 trunc mRNA was reduced compared to FL-WT mRNA (4.2 h vs 10.6 h) but was significantly increased compared to FL-W1282X mRNA (<<2 h).

**Figure 2.**
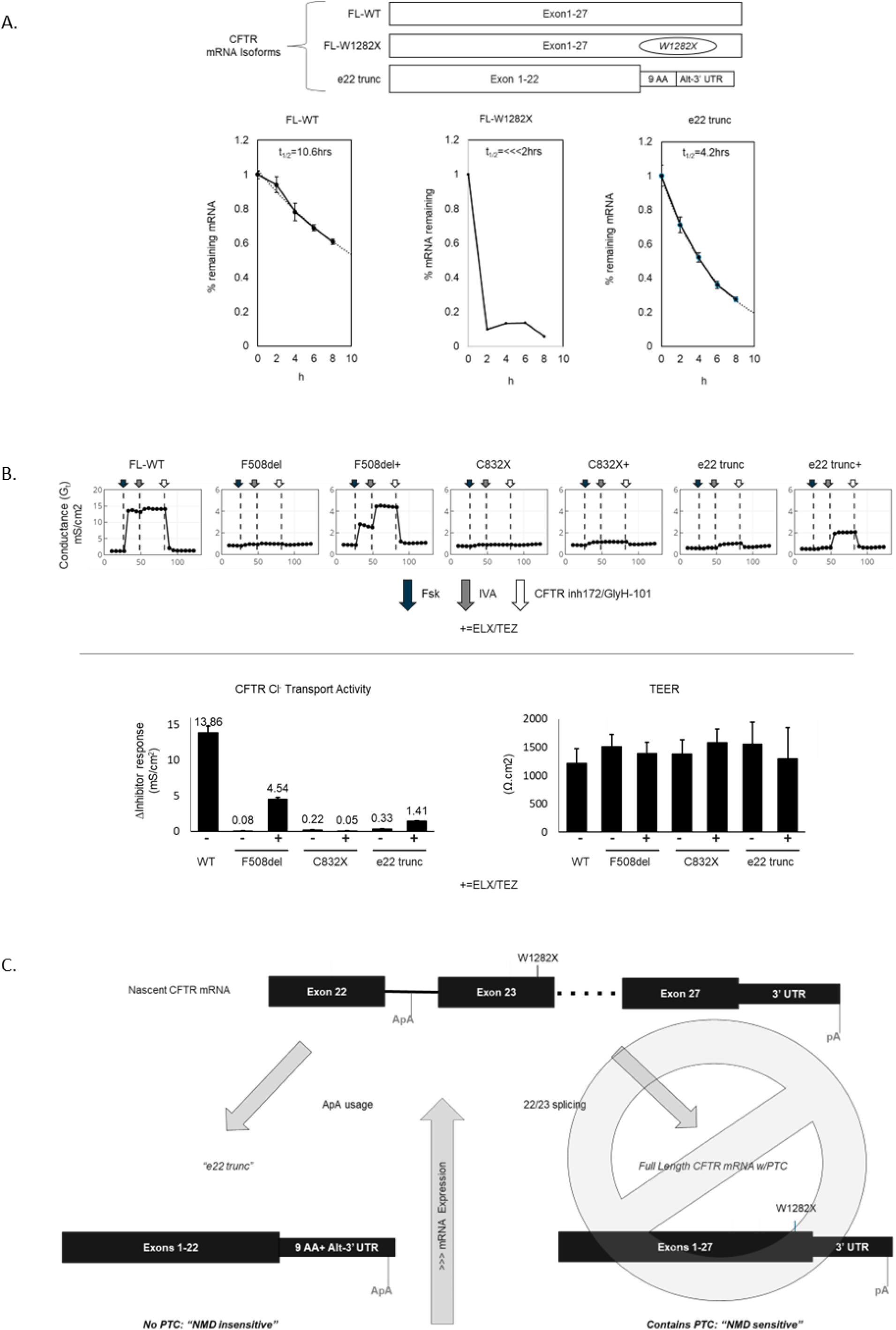
Therapeutic rationale for upregulation of a nonsense mediated decay (NMD) insensitive CFTR mRNA isoform for the treatment of 3’ CFTR PTC variants. **A.** mRNA stability of FL-WT, FL-W1282X, and e22 trunc. 16HBE14o- and 16HBEge W1282X cultures were treated at t=0 with Actinomycin D (5μg/ml) to block transcription and FL-WT, FL-W1282X, and e22 trunc mRNA were measured using ddPCR with specific probes targeting each mRNA isoform. % remaining mRNA was calculated relative to t=0 at 2,4,6,8, and 10 h and data was fit to an exponential decay function. **B**. Overexpression of WT *CFTR* (vehicle) and F508del, C832X and e22 trunc cDNAs in FRT cells with vehicle or 24 h pretreatment with 3/3 μM ELX/TEZ and 3 μM IVA acutely in TECC-24 assay. FRT parental, 96 h post transfection, treatment (apical and basolateral) 24 h prior to assay. CFTR Cl-transport activity as ΔInhibitor response (mS/cm^2^) calculated from plateau Fsk values. Transepithelial electrical resistance (TEER), a widely accepted measurement reporting on the integrity of tight junction in cell culture models of epithelial monolayers was unaffected by WT, F508del, C832X, or e22 trunc transfections or presence of 3/3 μM ELX/TEZ. **C**. Therapeutic rationale for exon 22/23 splice blocking to promote e22 trunc mRNA NMD escape and upregulation for 3’ terminal *CFTR* PTCs: Blockade of exon 22/23 splicing is predicted to increase nascent mRNA availability for intron 22 ApA usage and increase e22 trunc mRNA via NMD escape. pA=polyadenylation site and ApA=alternative polyadenylation site.

It is known that CFTR_trunc-1281_, missing most of NMD2 is trafficked to the apical membrane and retains partial channel function (9,10). To test whether e22 trunc protein also retains function, e22 trunc cDNA was expressed in FRT cells and CFTR channel function was evaluated. As illustrated in **Figure 2B**, cells transfected with e22 trunc cDNA and treated in assay with CFTR potentiator IVA exhibited a small increase in transepithelial conductance with forskolin that was further increased with pretreatment of CFTR correctors ELX/TEZ and reduced by CFTR inhibitor 172 (CFTR_inh_-172) and GlyH-101, responses consistent with e22 trunc channel activity. Transfection of FL-WT (positive control), F508del (ETI responsive CF variant) and C832X (ETI non-responsive negative control) were included for relative comparison. Given that e22 trunc mRNA has improved t_1/2_ compared to FL-W1282X and e22 trunc protein retains partial function, we asked if expression of e22 trunc could be modulated for therapeutic benefit of pwCF that harbor the W1282X variant. We hypothesized that inhibition of exon 22 pre-mRNA splicing would promote usage of intron 22 ApA sites and lead to upregulation of NMD insensitive e22 trunc mRNA and protein (**Figure 2C**). To test this, we gene-edited a 16HBE14o-cell line to remove the CFTR genomic sequence downstream of the eleven intron 22 ApA sites through ∼159 bp downstream of the 3’UTR, *16HBEge-ΔE23-3’UTR* (**Figure 3A**).

**Figure 3.**
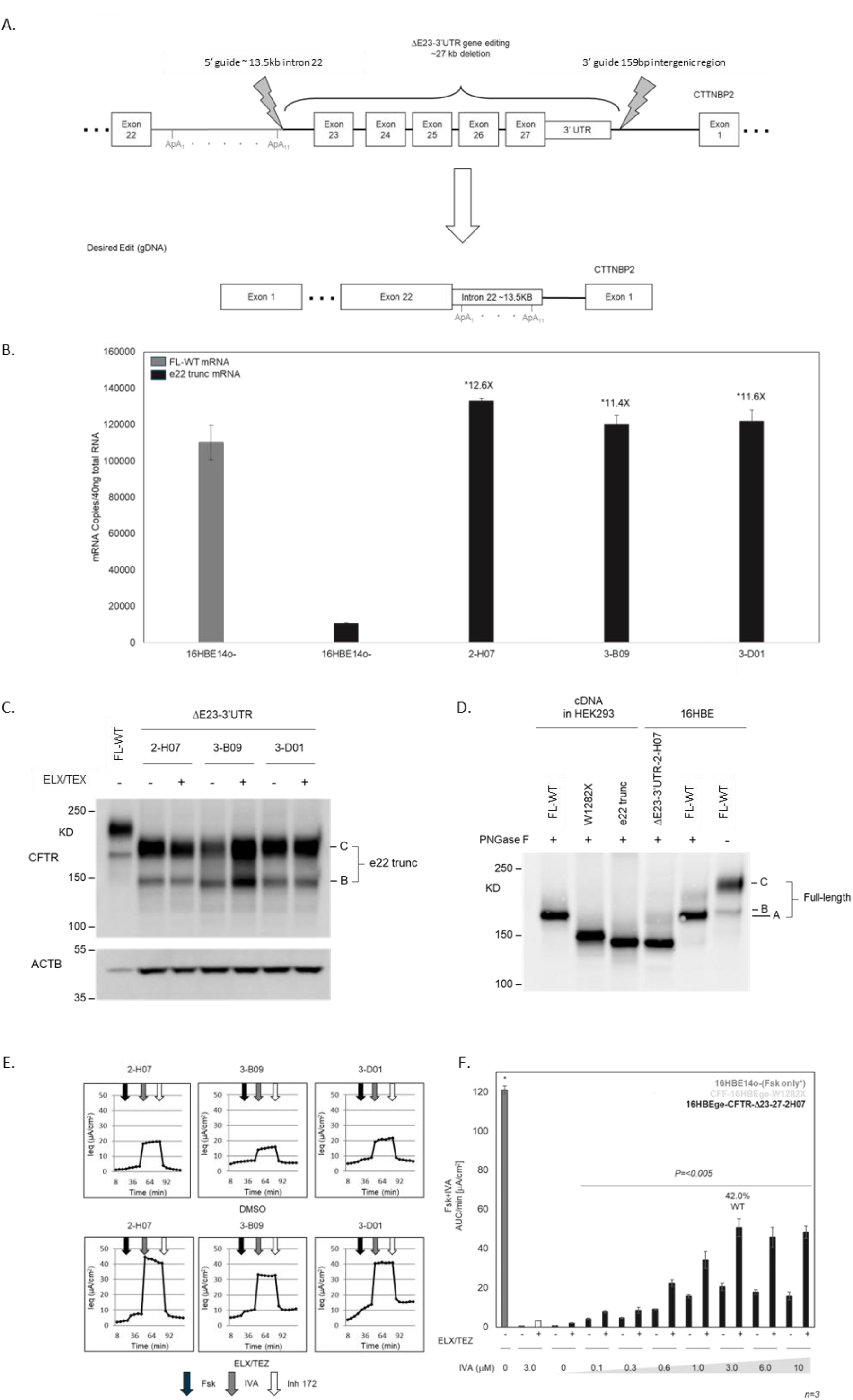
Abolishment of exon 22 splicing forces intron 22 ApA usage and elevates e22 trunc mRNA and protein levels that can be corrected with CFTR modulators to therapeutically relevant levels. **A.** Gene-edited cell model (ΔE23-3’UTR) to “force” e22 trunc mRNA expression in 16HBE14o-parental cells. **B**. ddPCR absolute mRNA copies of FL-WT Exon25/26 (black) and e22 trunc (grey) from 16HBE14o-parental cells and 16HBEge-ΔE23-3’UTR-clonal lines 2-H07, 3-B09, and 3-D01. **C**. Western blot analysis of 16HBE14o-parental line w/o ELX/TEZ and clones 2-H07, 3-B09, and 3-D01 ±ELX/TEZ. UNC 596 used for CFTR and ACTB used as loading control. **D**. Western blot of PNGaseF treated (de-glycosylated) FL-WT, FL-W1282X, and e22 trunc cDNA overexpression from HEK293 cells and 16HBEgeΔE23-3’UTR-2-H07 and 16HBE14o-parental cells. Samples were diluted to produce equivalent band intensity. **E**. Electrophysiological traces of 16HBEgeΔE23-3’UTR*-2-H07, 3-B09, and 3-D01* gene edited lines from TECC-24 assay after DMSO (vehicle) treatment (three upper panels) or 48 h pre-treatment with ELX/TEZ (3/3 μM) (three bottom panels). All samples treated in assay with IVA (1 μM). **F**. TECC-24 I_eq_ assay results of 16HBE14o-(grey bar) and 16HBEge W1282X (light grey bars) and IVA dose escalation in 16HBEgeΔE23-3’UTR*-2-H07* (black bars) ± ELX/TEZ 3/3 μM.

PCR screening was used to identify nine clones with the correct Δexon 23-3’UTR deletion junction. e22 trunc mRNA expression levels of the clones ranged from 0.5-1.2X of parental full-length CFTR mRNA levels (data not shown). Three *16HBEge-ΔE23-3’UTR* clones (2-H07, 3-B09, and 3-D01) with e22 trunc levels that were equivalent to parental WT-FL mRNA levels suggesting efficient usage of an intron 22 ApA site were selected for further characterization of CFTR mRNA, protein expression, and channel function. e22 trunc mRNA expression in the three clones increased ∼12-fold from parental e22 trunc levels (**Figure 3B**). Western blot analysis revealed the presence of appropriate molecular weight bands for partially (Band B) and fully (Band C) glycosylated e22 truncated CFTR protein (**Figure 3C and 3D**). As illustrated in **Figure 3E**, Fsk-stimulated CFTR channel function in the *16HBEge-ΔE23-3’UTR* clones was modest; however, pretreatment with CFTR correctors and acute exposure to a CFTR potentiator resulted in a ∼2 fold enhancement in channel function. Dose escalation of IVA with ELX/TEZ (3/3 μM) in the 2-H07 clone resulted in ∼42% WT function (**Figure 3F**). These results suggest that although e22 trunc protein has gating and potentially trafficking defects, CFTR modulators can significantly improve its channel function to therapeutically meaningful levels.

### ASO blockade of exon 22/23 splicing promotes intron 22 ApA usage and increases e22 trunc mRNA and protein levels which can be functionally rescued in the 16HBEge W1282X cell line model of CF

To extend our work to a therapeutically amenable approach, we applied ASOs to block exon 22/23 splicing and promote intron 22 ApA usage. Briefly, we designed 21 overlapping ASOs in 1 nucleotide “steps” across intron 22 splice donor (SD) (n=11) and splice acceptor (SA) sites (n=10) (**Supplementary Figure S2**). SD and SA ASOs were screened for the ability to increase e22 trunc mRNA in 16HBE14o-cells. Top performing SA-08 and SD-18 ASOs yielded 36.9% (**Supplementary Figure S3**) and 25.4% (**Supplementary Figure S4**) of FL-WT mRNA from 16HBE14o-cells, respectively, and were used for subsequent experiments.

The effect of ASO treatments on e22 trunc and FL-W1282X mRNA expression were assessed with ddPCR in 16HBEge W1282X cells (**Figure 4A**).

**Figure 4.**
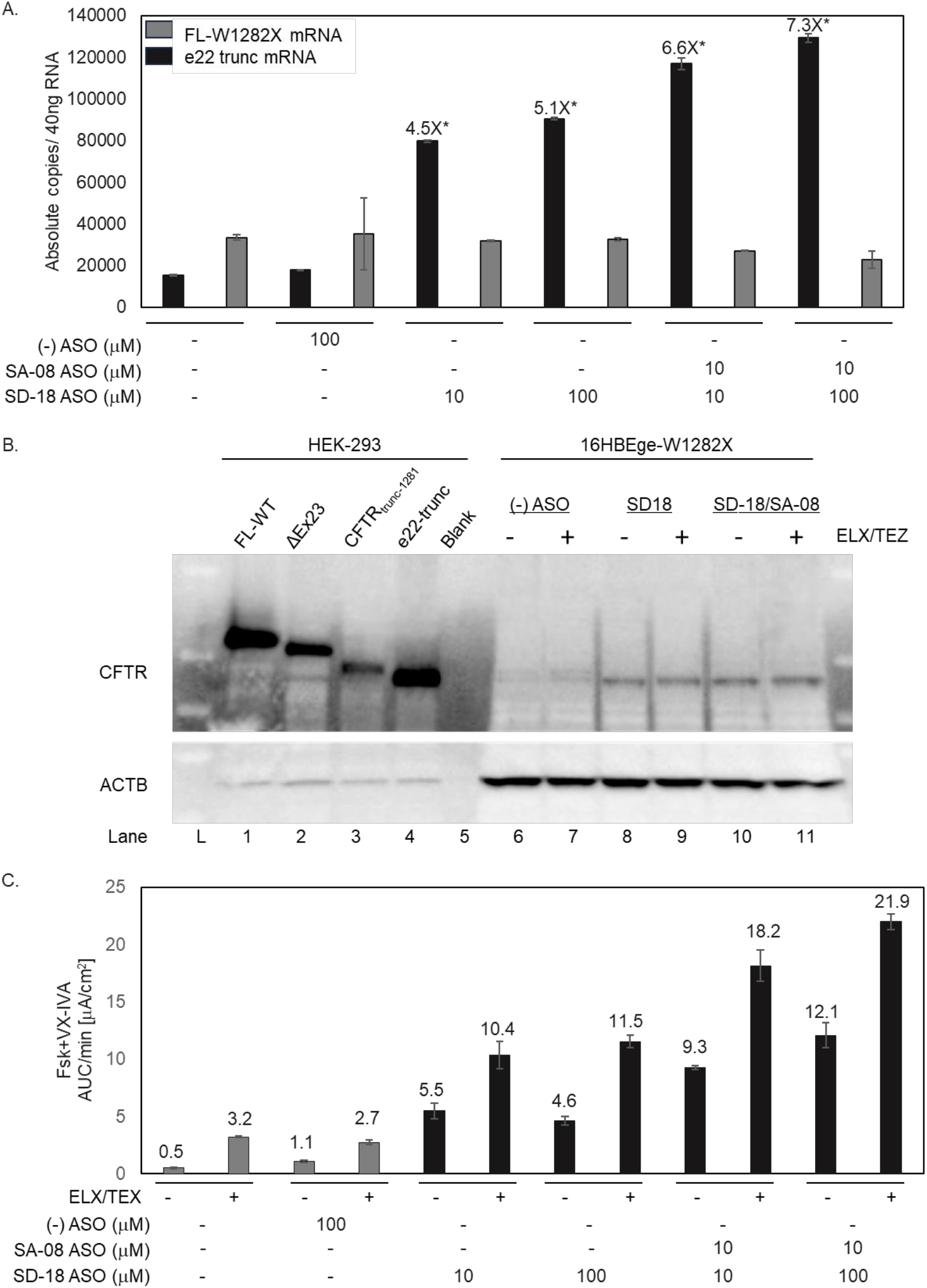
ASO blockade of exon 22/23 splicing promotes intron 22 ApA usage and increases e22 trunc mRNA and protein levels which can be functionally rescued in 16HBEge W1282X cell line model of CF. **A**. e22 trunc (black bars) and FL-W1282X (grey bars) mRNA quantitation via ddPCR as described above from 16HBEge W1282X post TECC-24 assay with DMSO, (-) ASO, SD-18, or SD-18/SA-08 48 h treatments and ± ELX/TEZ (3/3 μM) pretreatment for 48 h. All samples treated in assay with IVA (3 μM). **B**. Western blot analysis from 16HBEge W1282X cells post TECC-24 assay as described above. 16HBE14o- and *16HBEge-ΔE23-3’UTR* (ΔEx23), W1282X, and e22 trunc overexpressed in HEK293 cells as sizing controls (lanes 1-5). 16HBEge W1282X (-) ASO (10 μM), SD-18 (10 μM), and SD-18 (10 μM) & SA-08 (2 μM) pretreated for 48 h ± ELX/TEZ (3/3 μM) (lanes 6-11). CFTR UNC 596 used to detect CFTR and beta actin was used as loading control. All lysates were incubated with PNGaseF. Note the presence of W12821 (top band) and e22 trunc (lower band) in PBS control and the presence of only e22 trunc with ASO treatments. **C**. TECC-24 assay from 16HBEge W1282X with DMSO (vehicle) and (-) ASO (grey bars) or SD-18, SA-08, or SD-18 & SA-08 (black bars) 48 h treatments and ± ELX/TEZ (3/3 μM) pretreatment for 48 h. All samples treated in assay with IVA (3 μM).

FL-W1282X and e22 trunc mRNA were unchanged in (-) ASO (100 μM) control relative to vehicle control. (-) ASO targets the TNMD (tenomodulin) gene. TNMD is a cartilage-specific glycoprotein that was chosen because it is not expressed/detectable in 16HBE or hBE cultures. All SD-18 & SD-18/SA-08 combination treatments increased in e22 trunc mRNA levels relative to (-) ASO (100 μM): SD-18 (10 μM), 4.5 fold; SD-18 (100 μM), 5.1 fold; SD-18/SA-08 (10/10 μM), 6.6 fold; and SD-18/SA-08 (100/10 μM), 7.3 fold (*p=<0*.*05*). Unexpectedly, SD-18/ SA-08 (10/10 μM) resulted in a larger fold increase in e22 trunc mRNA than SD-18 (100 μM) treatment.

To evaluate the effect of ASO treatments on e22 trunc protein, 16HBE14ge W1282X cells were treated with (-) ASO (10 μM), SD-18 (10 μM) or SD-18/SA-08 (10/4 μM) ASOs for 48 h ± ELX/TEZ (3/3 μM) pretreatment. Western blots of CFTR protein from PNGaseF-deglycosylated cell lysates were compared to full length CFTR (∼170kDa, 1480 aa), CFTR Δexon23 (ΔEx23) (∼157 kDa, 1428 aa), CFTR_trunc-1281_ (∼141 kDa, 1281 aa), and e22 trunc (∼138 kDa, 1258 aa) HEK293 cDNA overexpression constructs (lanes 1-4). (**Figure 4B**). Consistent with the presence of e22 trunc and CFTR_trunc-1281_, faint bands aligning with e22 trunc (∼138 kDa) and CFTR_trunc-1281_ (∼141 kDa) overexpression controls were present in (-) ASO control cells independent of ELX/TEZ (3/3 μM) treatment (lanes 6 & 7). ASO treatments with SD-18 (10 μM) (lanes 8 & 9) or SD-18/SA-08 (10/4 μM) (lanes 10 & 11), ±ELX/TEZ (3/3 μM) incubation, resulted in a single high intensity band that aligned with the ∼138 kDa e22 trunc protein from the HEK293 overexpression control (lane 4). ELX/TEZ (3/3 μM, 48 h) treatment had minimal effect on intensity and no effect on the size of the protein products.

The TECC-24 assay (n=3) was then used to quantify the ASO-induced rescue of CFTR-mediated chloride transport in 16HBEge W1282X cells (**Figure 4C**). Treatment with (-) ASO (110 μM) 2.7±0.2 and 1.1±0.1 was approximately equivalent to vehicle alone 3.2±0.1 and 0.5±0.1 Fsk+IVA AUC/min (μA/cm^2^) ±ELX/TEZ (3/3 μM) respectively. All ASO treatments ±ELX/TEZ resulted in statistically significant increases in CFTR-mediated Cl-current relative to ASO control (*p<0*.*05*). SD-18 (10 μM), SD-18 (100 μM), SD-18/SA-08 (10/10 μM), and SD-18/SA-08 (100/10 μM) were 5.5±0.7, 4.6±0.4, 9.3±0.2, and 12.1±1.1 without ELX/TEZ and 10.4±1.2, 11.5±0.5, 18.2±1.3 and 21.9±0.7 Fsk+IVA AUC/min (μA/cm^2^) with ELX/TEZ (3/3 μM) respectively. As observed in the mRNA response data, SD-18/SA-08 (10/10 μM) resulted in a larger response than SD-18 (100 μM) treatment. A maximum functional response with SD-18/SA-08 (100/10 μM) + ELX/TEZ (3/3 μM) was equivalent to ∼20% WT.

### In primary cells homozygous for CFTR W1282X, ASO blockade of exon 22/23 splicing promotes intron 22 ApA usage resulting in increased expression of e22 trunc mRNA and rescue of CFTR function

To extend our analysis into a more translationally relevant model, we employed the “gold standard” CF model, hBE at ALI cultures. Unlike the previous 16HBEge W1282X cultures, hBEs at ALI form an airway-like epithelium that creates a barrier to free uptake. Therefore, to achieve sufficient upregulation of e22 trunc mRNA, we found it necessary to subject cultures to long ASO exposures starting at the undifferentiated state and refreshing ASOs with each media change throughout 21 days of differentiation. Undifferentiated *CFTR* W1282X^+/+^ hBE cells were treated with SD-18 (10 μM), SD-18 (100 μM), SD-18/SA-08 (10/10 μM), SD-18/SA-08 (100/10 mM), and (-) ASO (100 μM) ASOs for 48 h then differentiated for 21 days with media/ASO changes every two days. ASO treatment was discontinued 48 h prior to functional assays and cells were harvested to protein and mRNA quantification.

Primary *CFTR* W1282X^+/+^ hBE cells treated with (-) ASO (100 μM) had 495±42 and 715±36 baseline absolute copies/ 40ng total RNA of e22 trunc and FL-W1282X mRNA, respectively (**Figure 5A**).

**Figure 5.**
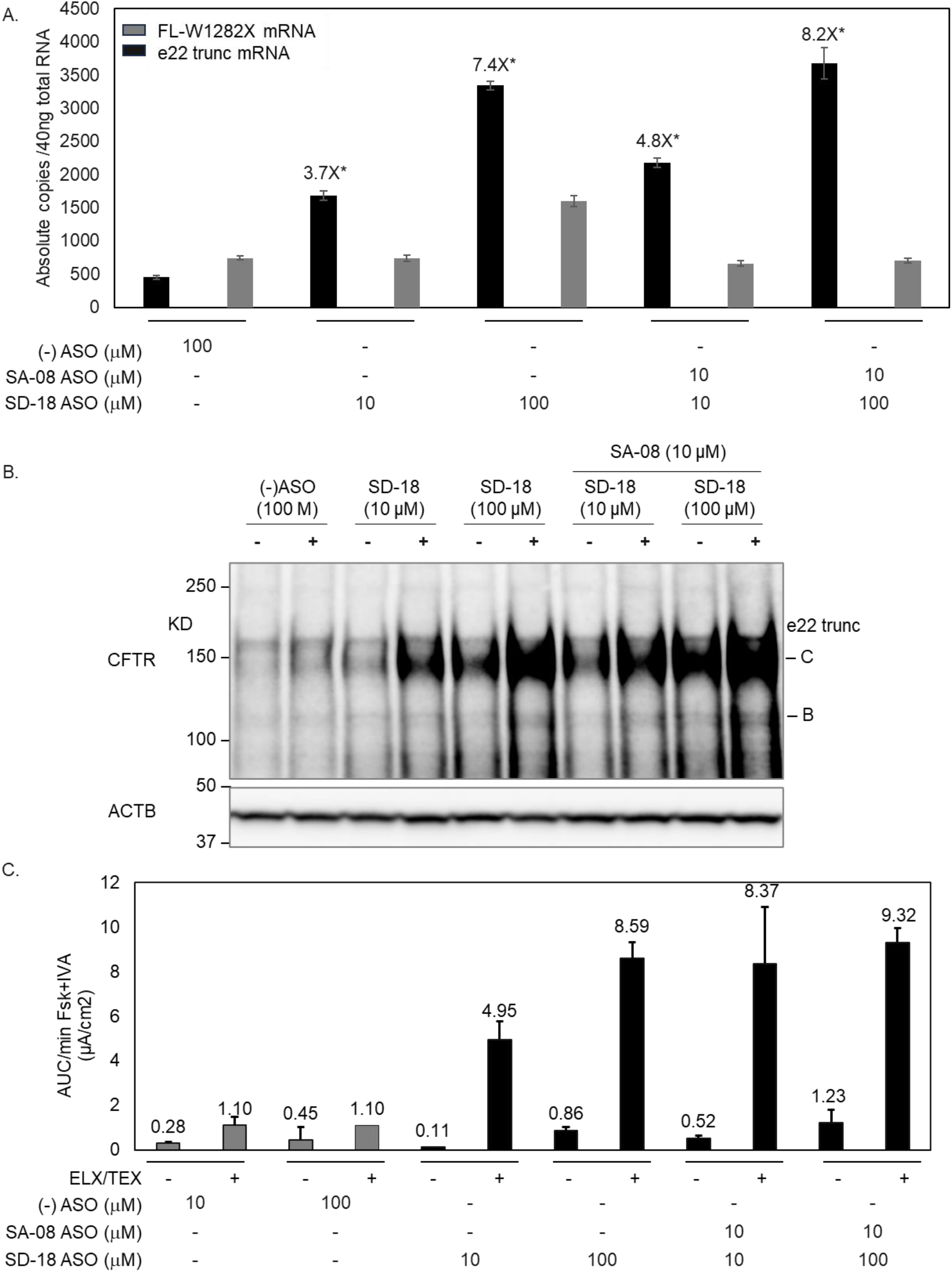
ASO blockade of exon 22/23 splicing promotes intron 22 ApA usage and increases e22 trunc mRNA and protein levels which can be functionally rescued in primary hBE cells homozygous for CFTR W1282X. **A**. FL-W1282X (grey bars) and e22 mRNA trunc quantitation (black bars) via ddPCR from fully differentiated *CFTR* W1282X^+/+^ hBE cells at ALI after TECC-24 assay. Cells were treated with (-) ASO, or SD-18, SA-08, or SD-18/SA-08 for 21 days and incubated ±ELX/TEZ (3/3 μM) for 48 h prior to the assay. All samples treated in assay with IVA (3 μM). **B**. Western blot analysis of fully ALI-differentiated *CFTR* W1282X^+/+^ hBE cells post TECC-24 assay as described above. Fully ALI-differentiated *CFTR* W1282X^+/+^ hBE cells untreated, or SD-18 (10 μM), or SD-18/SA-08 (10/2 μM) pretreated for 48 h ±ELX/TEZ (3/3 μM). CFTR detection with antibody UNC596; beta actin was used as loading control. Bar graph of CFTR Band C normalized to (-) ASO control without ELX/TEZ. **C**. TECC-24 assay from fully ALI-differentiated *CFTR* W1282X^+/+^ hBE cells treated with (-) ASO (grey bars) or SD-18, SA-08, or SD-18/SA-08 (black bars) for 21 days followed by ±) ELX/TEZ (3/3 μM) treatment for 48 h prior to the assay. All samples treated in assay with IVA (3 μM). Data represented as % WT (ratio of AUC/min W1282X (Fsk + IVA)/ WT (Fsk).

SD-18 (10 μM) increased e22 trunc 3.7-fold to 1683±65 absolute copies/40ng total RNA and FL-W1282X mRNA remained unchanged at 740±43 absolute copies/40ng total RNA. SD-18 (100 μM) further increased e22 trunc mRNA 7.4-fold to 3346±65 absolute copies/40ng total RNA and unexpectedly increased FL-W1282X mRNA to 1597±80 absolute copies/40ng total RNA. SD-18/SA-08 (10/10 μM) treatment increased e22 trunc mRNA 4.8-fold to 2176±72 absolute copies/40ng total RNA and again, FL-W1282X mRNA remained unchanged at 662±43 absolute copies/40ng total RNA. Lastly, consistent with the 16HBEge W1282X data, SD-18/SA-08 (100/10 μM) treatment resulted in the maximum induction of e22 trunc, 8.2-fold and 3681 ± 236 absolute copies/40ng total RNA. As expected, SD-18 (100 μM) & SA-08 (10 μM) treatment did not significantly impact FL-W1282X mRNA levels, 706 ±32 absolute copies/40ng total RNA.

Next, western analysis was used to measure the impact of ASOs on e22 trunc protein expression ±ELX/TEZ (3/3 μM) (**Figure 5B/Supplementary Figure S7**). Protein lysates were harvested from assay filters used in the TECC-24 functional assessments as described above. e22 trunc Band C intensity and ACTB bands were quantified, and Band C intensities were normalized to ACTB and referenced to (-) ASO (100 μM) ±ELX/TEZ (3/3 μM). All ASO blockade treatments increased Band C intensity relative to (-) ASO without ELX/TEZ. Treatment with ELX/TEZ further increased Band C intensity relative to vehicle control for all ASO treatments. As observed with e22 trunc mRNA copy number, the order of Band C intensity was SD-18/SA-08 (100/10 μM)> SD-18 (100 μM)> SD-18/SA-08 (10/10 μM)> SD-18 (10 μM), with relative intensities of 5.1, 4.9, 3.3, and 2.5 and 3.2, 2.4, 2.1, and 1.3 for with or without ELX/TEZ (3/3 μM), respectively.

To evaluate the functional impact of ASO treatments, TECC-24 assay were completed with fully differentiated *CFTR* W1282X^+/+^ hBE cells at ALI (**Figure 5C)**. (-) ASO (10 μM or 100 μM) (n=2) with vehicle resulted in negligible *CFTR*-mediated Cl^-^ current of 0.28±0.06 and 0.45±0.58 AUC/min Fsk+IVA (μA/cm^2^) that was increased marginally with ELX/TEZ (3/3 μM) to 1.1±0.37 and 1.1 (n=1) AUC/min Fsk+IVA (μA/cm^2^). Surprisingly, unlike the gene-edited model and the exon 22/23 splice blocking ASO treatments in 16HBEge W1282X, ASO treatments in hBEs at ALI in the absence of ELX/TEZ (3/3 μM) did not increase *CFTR*-mediated Cl^-^ currents even though increased mRNA and protein levels were observed. After 21-day ASO treatment at ALI followed by 48 h incubation with ELX/TEZ (3/3 μM), *CFTR*-mediated Cl^-^ currents were significantly increased (*p<0*.*05*) to 4.95±0.83, 8.59±0.74, 8.37±2.55, and 9.32±0.64 μA/cm^2^ with SD-18 (10 μM), SD-18 (100 μM), SD-18/SA-08 (10/10 μM), and SD-18/SA-08 (100/10 μM) relative to (-)ASO (100 μM), respectively. SD-18/SA-08 (100/10 μM) with ELX/TEZ (3/3 μM) resulted in ∼25% WT function.

### Conclusions

*CFTR* PTC variants are a challenging class of pathogenic variants to address therapeutically and there are currently no options available. Readthrough agents, such as aminoglycosides, that allow ribosomes to overread the PTC have been studied extensively since they in principle could provide a “universal therapeutic” for any disease caused by a nonsense variant (11,12,13,14,15). However, efficacy and potency are low, especially with respect to the level of functional restoration required for CF, the therapeutic index is narrow, and nephrotoxicity and ototoxicity are associated with chronic use (16,17). Additionally, the common notion that PTCs have a ∼10-fold higher propensity to be read through compared to natural termination codons (NTCs) (18,19,20,21) may be true at very low levels of readthrough but remains to be shown at a therapeutically meaningful range, calling into question the feasibility of global non-specific readthrough. Furthermore, protein products resulting from successful readthrough often contain a non-native amino acid that might negatively impact protein function and stability and thereby further limit the efficacy of this approach (10). Lastly, the problem of low mRNA template levels (10-20% of FL-WT mRNA) due to NMD remains a significant challenge that any readthrough therapeutic must overcome.

Here we describe an approach that could potentially address a subset of PTC variants that reside in the 3’ terminus of *CFTR* (e.g., W1282X, Q1313X, E1371X, etc.) for which there are no approved treatments. We first observed a lowly expressed e22 trunc mRNA, resulting from utilization of an alternative polyadenylation site in intron 22, in immortalized and primary cultures of human airway and intestinal epithelial cells. An important consequence of intron 22 ApA usage is that PTCs 3’ of exon 22 will be lost during mRNA processing of the nascent mRNA and will no longer trigger NMD, leading to a more stable transcript.

Previous studies have shown that a truncated CFTR protein (CFTR_trunc-1281_), missing most of NBD2 retains partial function that can be augmented by CFTR potentiator treatment. Heterologous expression of the e22 trunc cDNA in FRT cells and genetic deletion of exons 23-3’UTR in 16HBE14o-cells demonstrated that e22 trunc protein behaved similarly to the CFTR_trunc-1281_ protein. These unique features of e22 trunc mRNA and protein led us to hypothesize that modulation of e22 trunc mRNA/protein may have therapeutic potential for the treatment of CF caused by 3’ terminal PTCs in *CFTR*.

ASOs have recently garnered interest as a therapeutic modality for treatment of certain intractable human diseases. Notably, nusinersen, an ASO designed to promote inclusion of exon 7 in *SMN2* was approved for use in patients with Spinal Muscular Atrophy (SMA) (22). In vitro proof of concept (PoC) has shown that ASOs could provide therapeutic benefit to pwCF. These include ASOs targeting a cryptic splice site in intron 22 of the 3849+10kb C>T *CFTR* mutation (23), an ASO designed to induce exon 23 skipping for *CFTR* W1282X (24), and ASOs designed to prevent binding of exon junction complexes (EJC) downstream of exon 23 to attenuate NMD in a gene-specific manner for *CFTR* W1282X (25).

Our data demonstrate that steric blocking ASOs that target intron 22 splice donor and acceptor sites lead to upregulation of the e22 trunc mRNA, expression of truncated CFTR protein, and regulated CFTR channel function in non-functional W1282X immortalized airway epithelial and W1282X^+/+^ primary hBE ALI cell cultures treated with IVA/ELX/TEZ to ∼20% and ∼25% WT function respectively. These levels of functional rescue exceed the widely accepted belief that 10% improvement in an established *in-vitro* assay is predictive of clinical benefit across a cohort of pwCF (26).

This approach would be applicable to pwCF that have variants that reside in exons 23-27 such as: 1) PTCs, 2) indels that result in in-frame PTCs, or 3) loss of function variants that are non-responsive to available highly effective modulator therapy (HEMT). However, we would not expect this approach to be appropriate for pwCF that have 1 or more HEMT-responsive alleles. Additionally, while the ASO targeting the 3849+10kb C>T variant, restores FL-WT mRNA, our data suggest that upregulation of e22 trunc protein will require co-administration of CFTR modulators, at minimum a potentiator, to provide therapeutic benefit.

Given the limited success in development of therapies for pwCF harboring PTC variants, novel and/or combination approaches may be required for their treatment. Here we show that inhibition of splicing via gene editing or ASO blockage increases e22 trunc mRNA, protein, and function to the range that is expected to provide therapeutic benefit. We speculate that further improvement of functional response may be achieved by improving e22 trunc mRNA t_1/2_ by promoting usage of downstream ApA site and increasing the 3’ alternative UTR length. Additionally, next generation potentiators could also increase the magnitude of functional restoration achievable with the e22 trunc protein. This work provides a foundation for further development of a promising approach for the treatment of pwCF harboring 3’ terminal PTCs.

## Abbreviations

CF: cystic fibrosis
CFTR: cystic fibrosis transmembrane conductance regulator
pwCF: people with cystic fibrosis
e22 trunc mRNA: CFTR Exon 22 truncated mRNA
ApA: alternative polyadenylation
ASO: antisense oligonucleotide
2’-MOE: 2-O-Methyethyl
PS: phosphorothioate
PNGaseF: peptide-N-glycosidase F
Fsk: forskolin
ELX: elexacaftor
TEZ: tezacaftor
IVA: ivacaftor
ETI: elexacaftor/tezacaftor/ivacaftor
WT: wild type
FRT cells: Fischer rat thyroid cells
G_t_: transepithelial conductance
I_eq_: equivalent short-circuit current
CFTR_inh_-172: CFTR inhibitor 172
TEER: Transepithelial electrical resistance
IO: intestinal organoid
UNC 596: CFTR detection antibody

## Acknowledgments

Martin Mense, Calvin Cotton, John Carulli for scientific discussion and careful review of manuscript.

Michelle Hasting for discussions about delivery options for ASO.

Garry Cutting & Karen Raraigh for assessment of fraction of pwCF that harbor variants that may be amenable to e22 trunc therapy.

## Author contributions

NA & JY conceived of the original concept of modulating ApA usage for therapeutic benefit of 3’ PTCs. HB conceived of gene edited model to force intron 22 ApA usage. HV designed guide RNAs for gene edited model. NA, MA, and JY designed, analyzed, and interpreted experiments. MA, JY, PB, JH, YC, and CM carried out experiments. AS gave input for statistical analysis. NA wrote the manuscript. MM and CC provided critical feedback and edited the manuscript.

## This work is subject of the following patent application

International Application No. PCT/US2023/075747 Priority: US 63/412771 filed October 3, 2022

Title: COMPOSITIONS AND METHODS FOR MODULATION OF CFTR

Applicant: Cystic Fibrosis Foundation

Inventors: Normand E. Allaire, Jae Seok YOON, Mathew Armstrong

