## Supplemental figures for "Upregulation of a nonsense mediated decay (NMD) insensitive CFTR mRNA isoform has therapeutic potential for the treatment of 3’ CFTR PTC variants"

### Supplementary data: Table S1: Sequences of intron 22 splice donor, acceptor, and control ASOs

| Blocking ASO Name | Target Sequence | Blocking ASO-Reverse Complement of Target | Start | End | Chrom | SPAN | Reference Genome |
|---|---|---|---|---|---|---|---|
| SA-01 | TCACTTTTACCTTATAGGTGGG | CCCACCTATAAGGTAAAAGTGA | 117642421 | 117642442 | Chr7 | 22 | GRCh38/hg38 |
| SA-02 | ATCACTTTTACCTTATAGGTGG | CCACCTATAAGGTAAAAGTGAT | 117642420 | 117642441 | Chr7 | 22 | GRCh38/hg38 |
| SA-03 | CATCACTTTTACCTTATAGGTG | CACCTATAAGGTAAAAGTGATG | 117642419 | 117642440 | Chr7 | 22 | GRCh38/hg38 |
| SA-04 | CCATCACTTTTACCTTATAGGT | ACCTATAAGGTAAAAGTGATGG | 117642418 | 117642439 | Chr7 | 22 | GRCh38/hg38 |
| SA-05 | CCCATCACTTTTACCTTATAGG | CCTATAAGGTAAAAGTGATGGG | 117642417 | 117642438 | Chr7 | 22 | GRCh38/hg38 |
| SA-06 | TCCCATCACTTTTACCTTATAG | CTATAAGGTAAAAGTGATGGGA | 117642416 | 117642437 | Chr7 | 22 | GRCh38/hg38 |
| SA-07 | ATCCCATCACTTTTACCTTATA | TATAAGGTAAAAGTGATGGGAT | 117642415 | 117642436 | Chr7 | 22 | GRCh38/hg38 |
| SA-08 | GATCCCATCACTTTTACCTTAT | ATAAGGTAAAAGTGATGGGATC | 117642414 | 117642435 | Chr7 | 22 | GRCh38/hg38 |
| SA-09 | TGATCCCATCACTTTTACCTTA | TAAGGTAAAAGTGATGGGATCA | 117642413 | 117642434 | Chr7 | 22 | GRCh38/hg38 |
| SA-10 | GTGATCCCATCACTTTTACCTT | AAGGTAAAAGTGATGGGATCAC | 117642412 | 117642433 | Chr7 | 22 | GRCh38/hg38 |
| SD-06 | CAGAGGGTGAGATTTGAACACT | AGTGTTCAAATCTCACCCTCTG | 117627765 | 117627786 | Chr7 | 22 | GRCh38/hg38 |
| SD-11 | AGAGGGTGAGATTTGAACACTG | CAGTGTTCAAATCTCACCCTCT | 117627766 | 117627787 | Chr7 | 22 | GRCh38/hg38 |
| SD-12 | GAGGGTGAGATTTGAACACTGC | GCAGTGTTCAAATCTCACCCTC | 117627767 | 117627788 | Chr7 | 22 | GRCh38/hg38 |
| SD-13 | AGGGTGAGATTTGAACACTGCT | AGCAGTGTTCAAATCTCACCCT | 117627768 | 117627789 | Chr7 | 22 | GRCh38/hg38 |
| SD-14 | GGGTGAGATTTGAACACTGCTT | AAGCAGTGTTCAAATCTCACCC | 117627769 | 117627790 | Chr7 | 22 | GRCh38/hg38 |
| SD-07 | GGTGAGATTTGAACACTGCTTG | CAAGCAGTGTTCAAATCTCACC | 117627770 | 117627791 | Chr7 | 22 | GRCh38/hg38 |
| SD-15 | GTGAGATTTGAACACTGCTTGC | GCAAGCAGTGTTCAAATCTCAC | 117627771 | 117627792 | Chr7 | 22 | GRCh38/hg38 |
| SD-16 | TGAGATTTGAACACTGCTTGCT | AGCAAGCAGTGTTCAAATCTCA | 117627772 | 117627793 | Chr7 | 22 | GRCh38/hg38 |
| SD-17 | GAGATTTGAACACTGCTTGCTT | AAGCAAGCAGTGTTCAAATCTC | 117627773 | 117627794 | Chr7 | 22 | GRCh38/hg38 |
| SD-18 | AGATTTGAACACTGCTTGCTTT | AAAGCAAGCAGTGTTCAAATCT | 117627774 | 117627795 | Chr7 | 22 | GRCh38/hg38 |
| SD-08 | GATTTGAACACTGCTTGCTTTG | CAAAGCAAGCAGTGTTCAAATC | 117627775 | 117627796 | Chr7 | 22 | GRCh38/hg38 |
| CEP290 | CATGAAGGTCTTCCTCATGC | GCATGAGGAAGACCTTCATG | 100542440 | 100542459 | Chr10 | 20 | GRCm38/mm10 |
| Scrambled | N/A | CTTCCCTGAAGGTTCCTC | N/A | N/A | N/A | 18 | N/A |
| TNMD | AAAGGGATTGAACAAAATGAAC | GTTCATTTTGTTCAATCCCTTT | 100599078 | 100599099 | ChrX | 22 | GRCh38/hg38 |

***Table S1.*** Splice donor and acceptor ASOs named with prefix SD and SA respectively.

### Supplementary data: Table S2: Sequences of ddPCR assays

| Assay | CFTR E22 trunc ddPCR assay (85bp amplicon) | CFTR Exon25/26 ddPCR assay (123 bp amplicon) |
|---|---|---|
| Forward Primer | ACAGAAGGTGGAAATGCCATA | TTGCTGCTTGATGAACCCA |
| Reverse Primer | AAGCAAGCAGTGTTCAAATCTC | CATTGCTTCTATCCTGTGTTCAC |
| Probe | /56-FAM/TCTCAATAA/ZEN/GTCCTGGCCAGAGGGT/3IABkFQ/ | /56-FAM/TGCTTGTTT/ZEN/TAGAGTTCTTCTAATTATTTGGTATGTTACT/3IABkFQ/ |

***Table S2.*** CFTR E22 trunc ddPCR assay used as a proxy of exon 22 truncated mRNA and CFTR Exon 25/26 ddPCR assay used as a proxy of full length CFTR mRNA

### Supplementary data: Figure S1. Genomic evidence of intron 22 alternatively polyadenylated transcripts

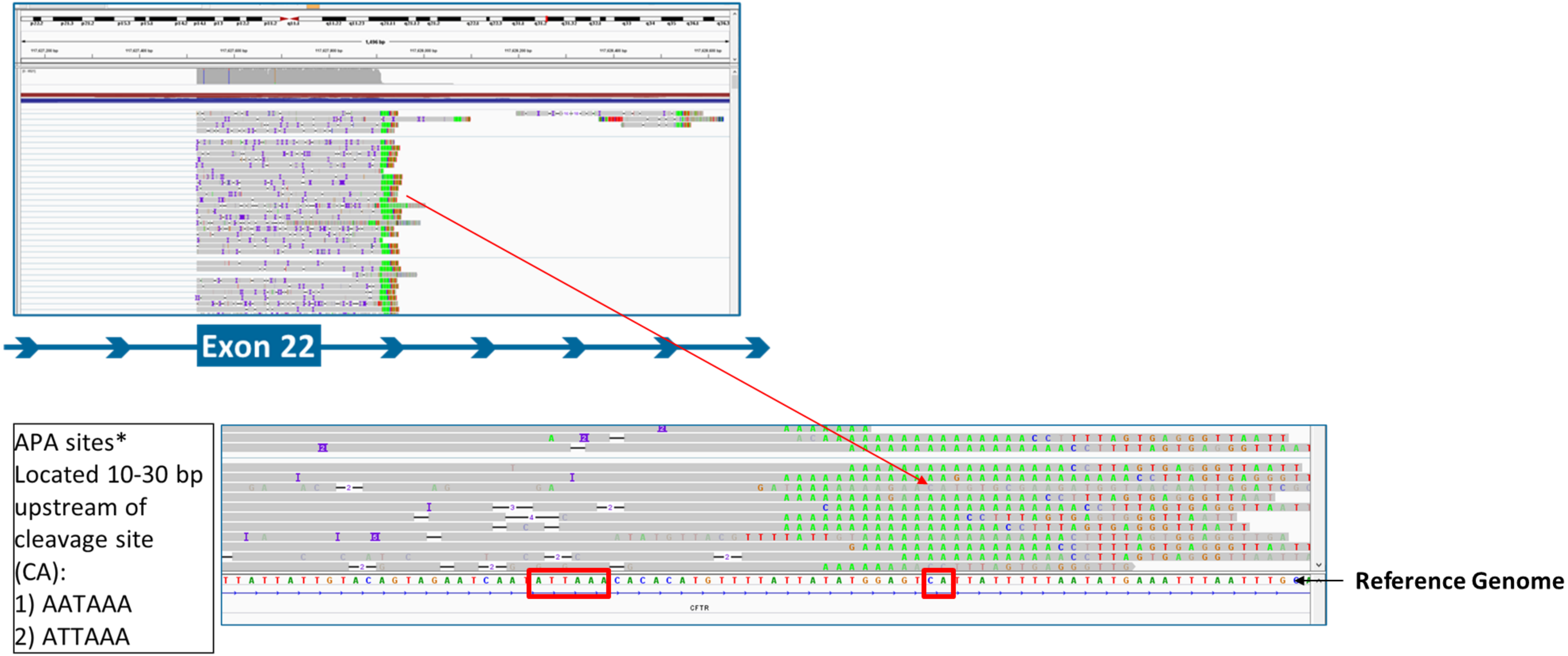


**Figure S2.** Integrated Genomic Viewer (IGV) image showing extension of mRNA reads into intron 22. Green adenine residues that do not align to genomic reference provide evidence of post transcriptional polyadenylation. Red boxes indicate alternative polyadenylation consensus sequence and dinucleotide cleavage site.

### Supplementary data: Table S3: e22 trunc and full length CFTR mRNA levels in wild type and W1282X+/+ primary airway, intestinal, and immortalized cells.

| Cell Type | Subject | Genotype | Exon 25/26- % FL CFTR +/- STD | e22 trunc- % FL CFTR +/- STD |
|---|---|---|---|---|
| hBE @ALI | Subject 1 | WT | 83.8 +/-8.9 | 9.3 +/-1.2 |
| hBE @ALI | Subject 2 | WT | 116.2 +/-3.6 | 6.5 +/-4.1 |
| hBE @ALI | Subject 3 | W1282X+/+ | 8.0 +/-2.4 | 5.7 +/-2.4 |
| IO | Subject 4 | WT | 100.0 +/-8.2 | 6.9 +/-2.1 |
| IO | Subject 5 | W1282X+/+ | 11.3 +/-2.5 | 7.1 +/-1.8 |
| IO | Subject 6 | W1282X+/+ | 19.9 +/-1.0 | 11.6 +/-1.7 |
| 16HBE | 16HBE14o- | WT | 100.0 +/-8.7 | 8.6 +/-2.9 |
| 16HBE | CFF-16HBEge-CFTR-W1282X | W1282X+/+ | 18.5 +/-1.7 | 10.4 +/-0.9 |

**Table S3.** RNA was harvested from wild type and W1282X$^{+/+}$ IOs, hBE at ALI, and 16HBE14o- and 16HBEge-W1282X cells. Fractions of e22 trunc and full length CFTR mRNA relative to WT full length CFTR mRNA +/- standard deviation (STD) were calculated from ddPCR values using CFTR e22 trunc and CFTR Exon 25/26 assays.

### Supplementary data: Figures S2-S5. Development and validation of ASOs block exon 22/23 splicing, induce intron 22 ApA usage, and exon 22 trunc mRNA upregulation

***Figure S1.*** 2’MOE ASO (22 mers) were tiled in 1 nucleotide “steps” (5’>3’) across intron 22 splice donor (n=11) beginning at HG38 chr 7: 117,627,765 and 1 nucleotide “steps” (3’>5’) across splice acceptor sites (n=10) beginning at HG38 chr 7: 117,642,442. ***Figure S2.*** 16HBE14o- cells were treated with Intron 22 splice donor ASOs (10μM) (n=11) and scrambled (10μM) CEP290 (10μM), or untreated for 48hrs. ddPCR of absolute levels of exon 22 truncated CFTR mRNA was assayed from 40ng total RNA. Data is expressed as % full length CFTR from untreated 16HBE14o- WT cells. ***Figure S3.*** 16HBE14o- cells were treated with Intron 22 splice acceptors ASOs (10μM) (n=10) and scrambled (10μM), CEP290 (10uM), or untreated for 48hrs. ddPCR of absolute levels of exon 22 truncated CFTR mRNA was assayed from 40ng total RNA. Data is expressed as % full length CFTR from untreated 16HBE14o- WT cells. ***Figure S4.*** Locations of intron 22 splice donor blocking ASOs. Top hit I22_SD_SBO_18 grey box. ***Figure S5.*** Locations of intron 22 splice acceptor blocking ASOs. Top hit I22_SA_SBO_08 grey box.

### Supplementary data: Figure S2. Design of intron 22 Splice Acceptor ASO blockage using a 1 NT walk

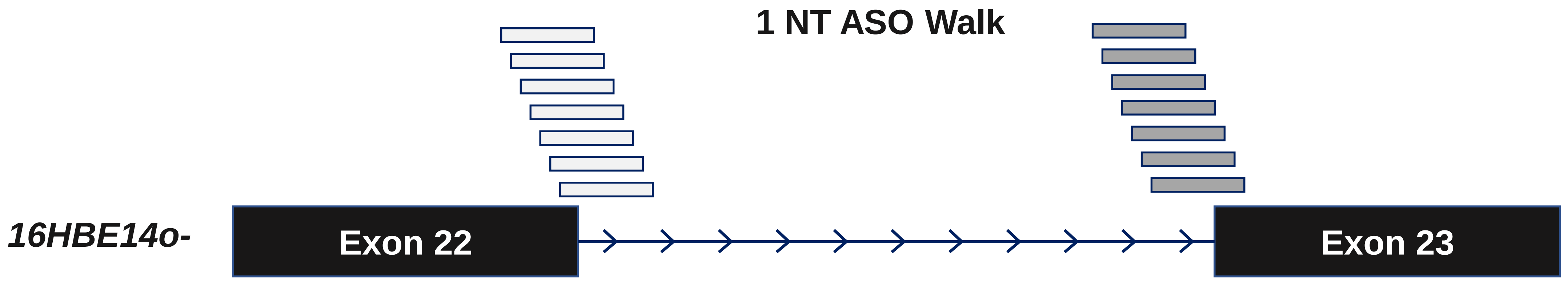

### Supplementary data: Figure S3. Splice Donor 1NT Walk (48hrs @ 10uM)

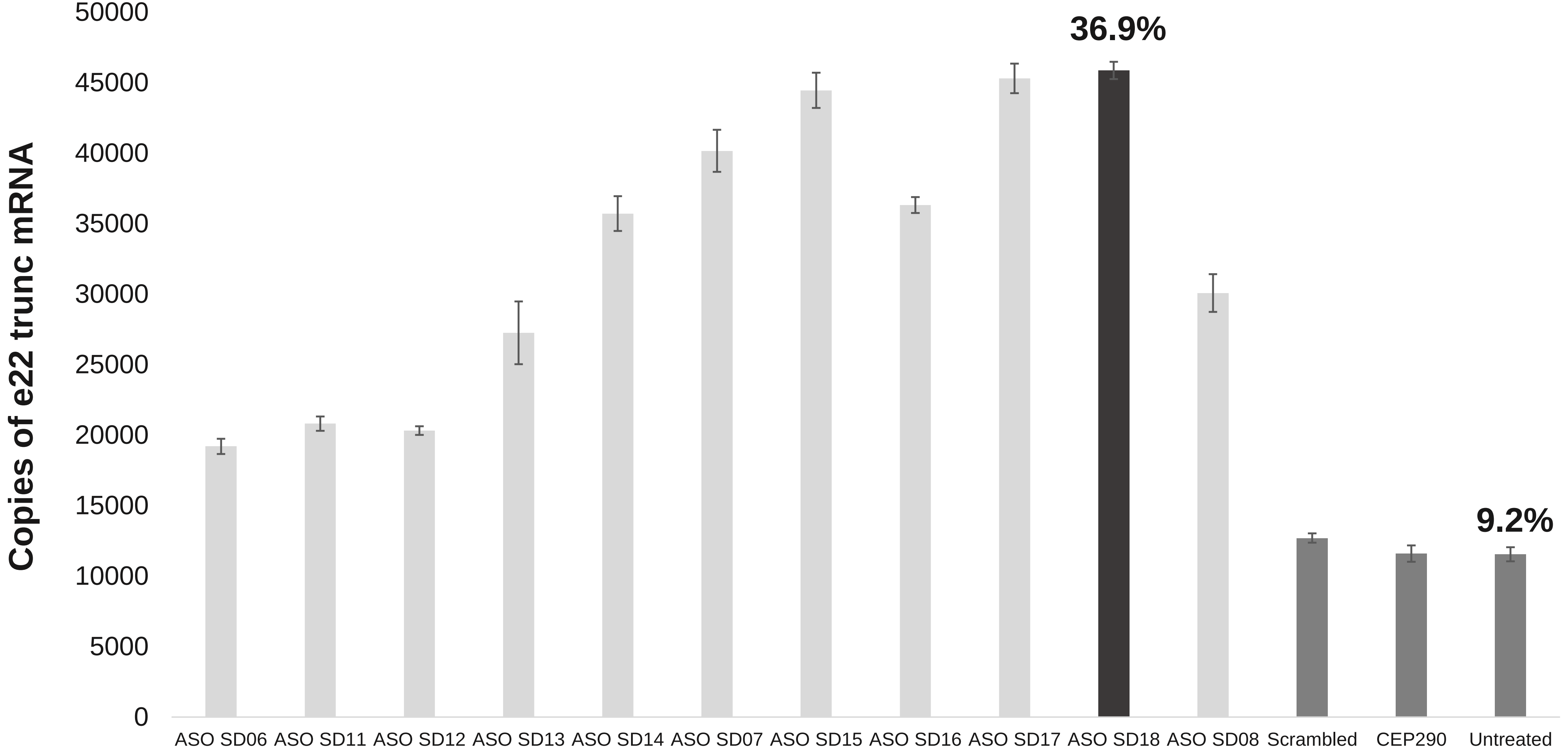


***% of FL WT CFTR (~125K copies)**

### Supplementary data: Figure S4. Splice Acceptor 1NT Walk (48hrs @ 10uM)

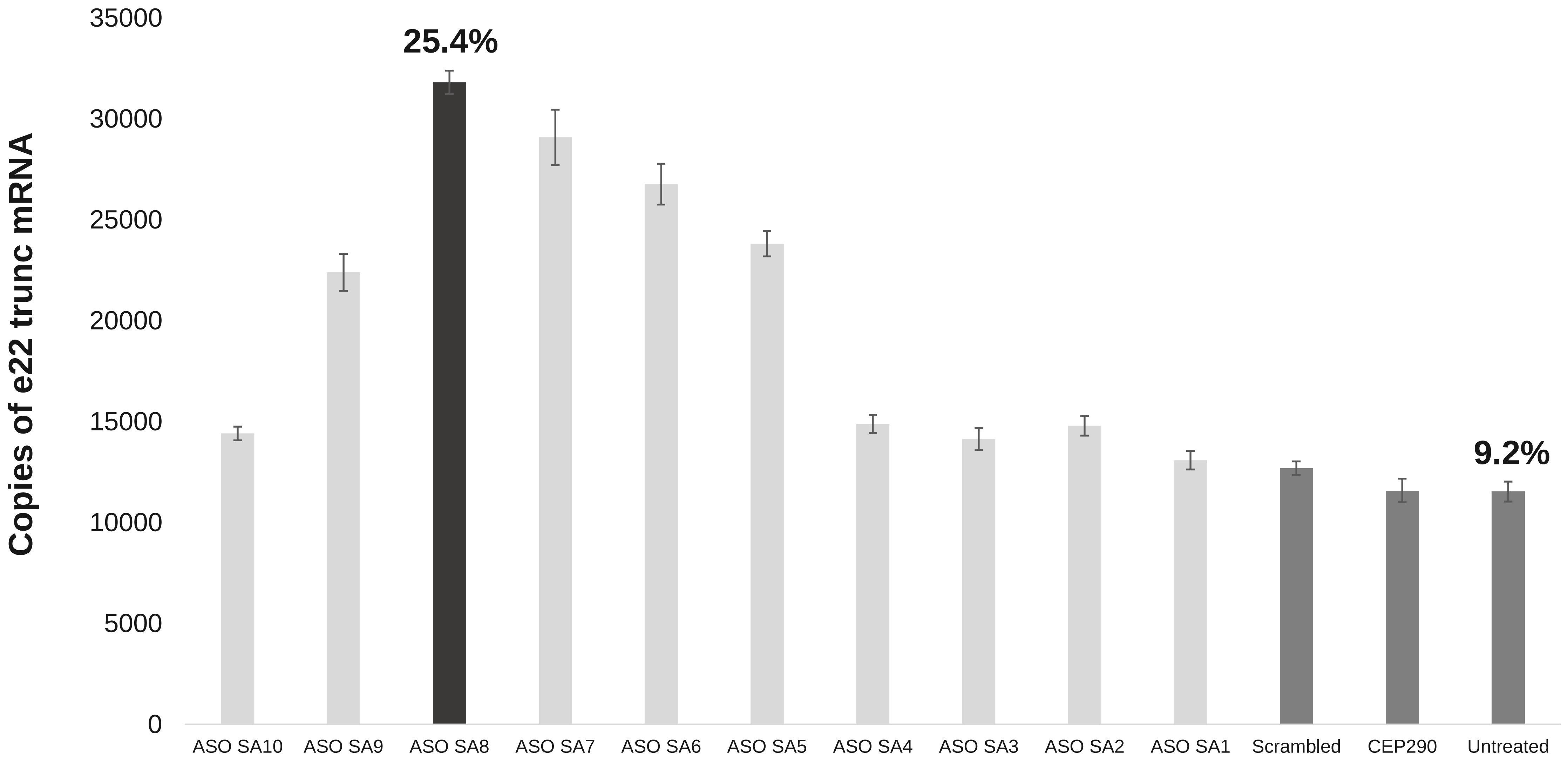


*% of FL WT CFTR (~125K copies)

### Supplementary data: Figure S5. Top SD Hit- i22_SD_SBO_18 (SD-18)

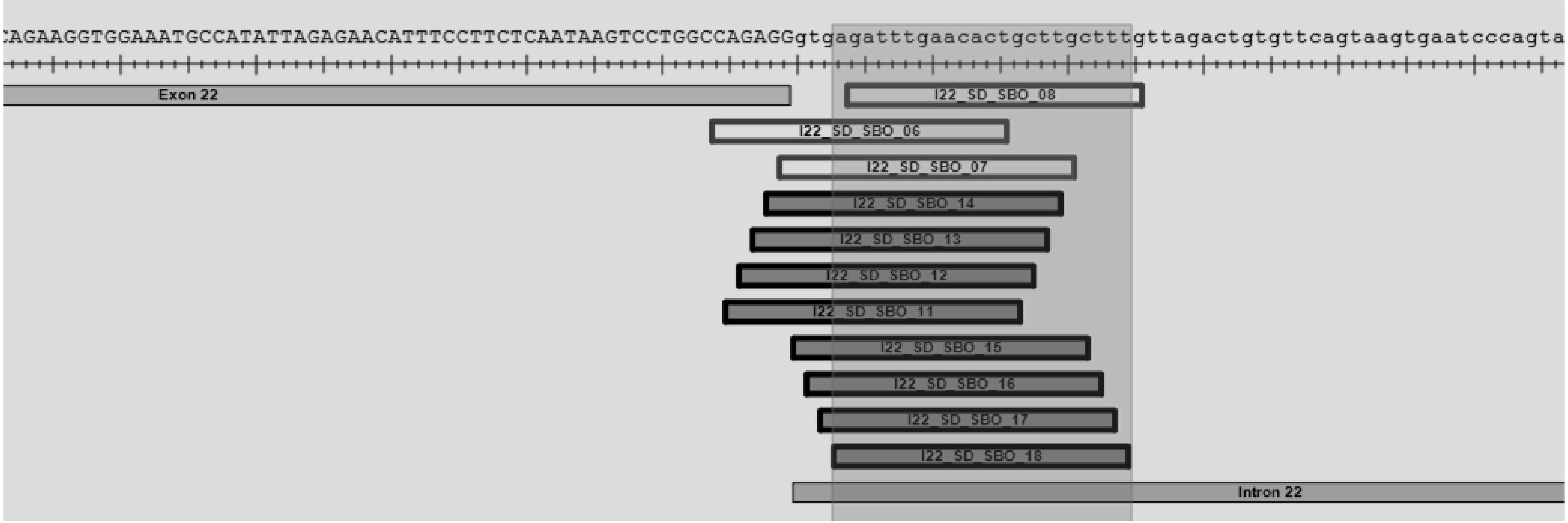

### Supplementary data: Figure S6. Top SA Hit- i22_SA_SBO_08 (SA-08)

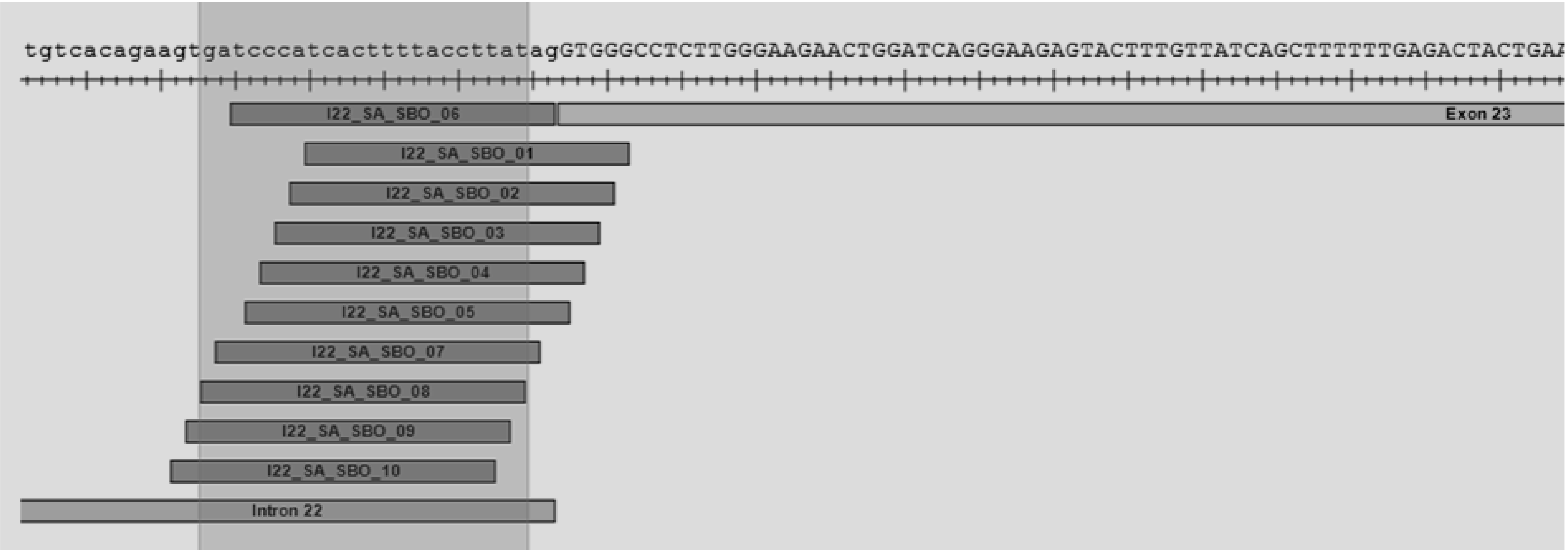

### Supplementary data: Figure S7. Quantitation of western blot CFTR Band C intensities from fully differentiated hBE W1282X$^{+/+}$ treated with ASOs

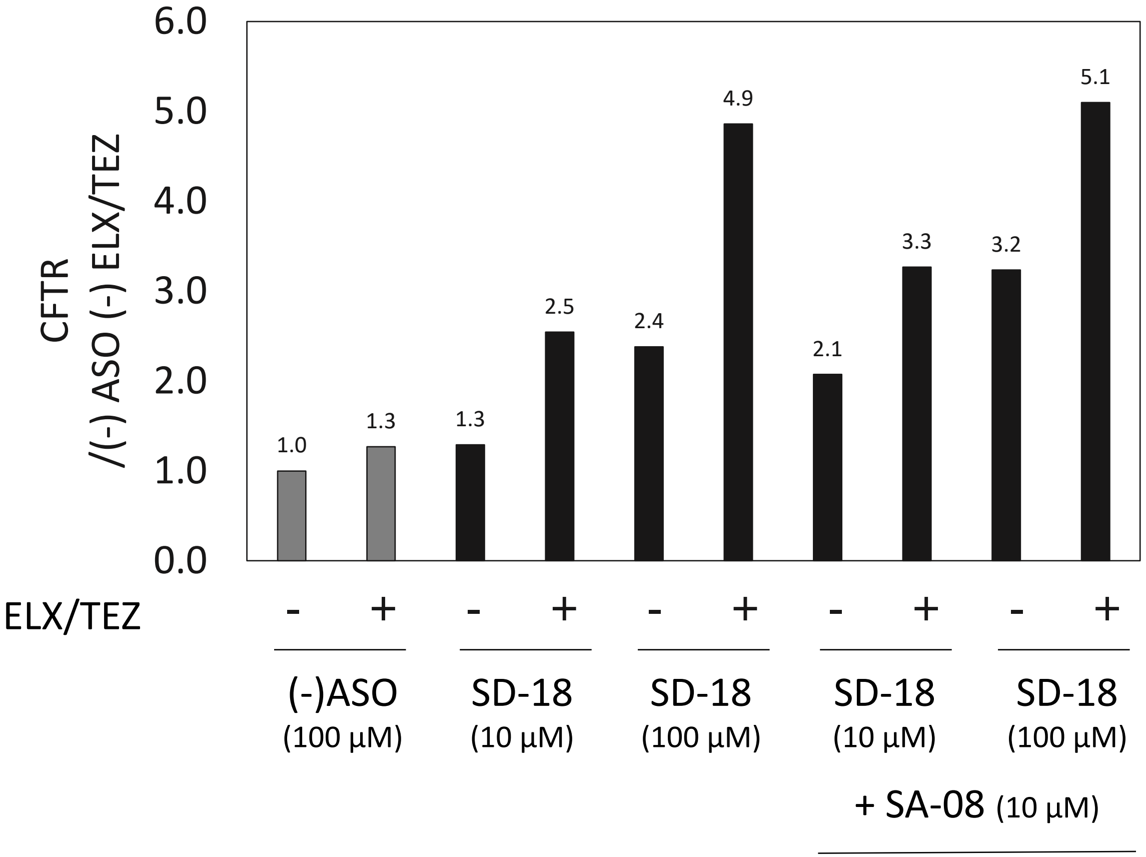


***Figure S7.*** e22 trunc Band C intensity and ACTB band were quantified using densitometry. Grey bars are (-) ASO targeting TNMD gene which is not expressed in primary airway cultures.
